# Ancestral resurrection reveals mechanisms of kinase regulatory evolution

**DOI:** 10.1101/331637

**Authors:** Dajun Sang, Sudarshan Pinglay, Sezen Vatansever, Hua Jane Lou, Benjamin Turk, Zeynep H. Gümüş, Liam J. Holt

## Abstract

Protein kinases are crucial to coordinate cellular decisions and therefore their activities are strictly regulated. We used ancestral resurrection to uncover a mechanism underlying the evolution of kinase control within the ERK family of Mitogen Activated Protein Kinases (MAPKs). Kinase activities switched from high to low intrinsic autophosphorylation at the transition from the ancestors of ERKs1-5 and ERKs1-2. A shortening of the loop between β3-αC and a mutation in the gatekeeper residue drove this transition. Molecular dynamics simulations suggested that the change in the β3-αC loop length affected kinase cis-autophosphorylation by altering the positioning of catalytic residues and by allowing greater flexibility in the L16 kinase loop. This latter effect likely synergizes with the known role of gatekeeper mutations in facilitating domain closure and thus kinase activation, providing a rationale for the synergy between the two evolutionary mutations. Our results shed light on the evolutionary mechanisms that led to tight regulation of a central kinase in development and disease.

## Introduction

Protein kinases regulate most aspects of cell life, including growth, differentiation, proliferation and apoptosis. They are one of the largest protein families in eukaryotes (Lahiry, et al. 2010) with more than 500 kinases in the human genome, comprising 2% of all protein coding genes. Due to their centrality in signaling networks, kinase activity has evolved to be precisely regulated by a myriad of distinct mechanisms. A breakdown in kinase regulation is the underlying cause of many human diseases; in particular, kinase mutations drive many forms of cancer.

Kinases have evolved a multitude of mechanisms to control their activation. Some examples such as the requirement for allosteric modulators, regulation by inhibitory interactions, and localization control are highly variable across the kinome. However, the activation of most kinases requires phosphorylation of the activation loop, a conserved structural feature that connects the N and C-terminal lobes of the kinase domain. Phosphorylation of a conserved TxY motif within this loop affects both substrate binding and the arrangement of catalytic residues (Johnson, et al. 1996; Nolen, et al. 2004), leading to robust regulation of enzyme activity.

Extensive structural and biochemical studies have afforded substantial insight into the mechanisms of kinase regulation by activation loop phosphorylation. In the mitogen activated protein kinases (MAPKs) ERK1 and ERK2, several structural elements have been shown to be important for activity. These include the P+1 binding pocket that binds the substrate peptide proline, the regulatory αC helix, and the L16 loop. Upon phosphorylation of the TxY threonine and tyrosine residues, a number of changes occur. A network of electrostatic contacts with the phospho-threonine and phospho-tyrosine leads to a conformation change around the P+1 substrate binding site, enabling substrate binding. The regulatory αC helix rotates inwards, allowing a glutamine on the αC helix (Glu-69) to form a salt bridge with Lysine (Lys-52) on the β3 strand that is crucial for catalytic activity. Extension of the C terminal L16 loop enables rotation of the N- and C-terminal domains, leading to “domain closure”, bringing Lys-52 closer to other catalytic residues, such as aspartates in the DFG and HRD motifs. Together, these conformational changes lead to up to 1000 fold increase in activity (Zhang, et al. 1994; Canagarajah, et al. 1997; Roskoski 2012).

Some kinases are able to autophosphorylate and thus autoactivate. Multiple mechanisms for autophosphorylation have been proposed (Beenstock, et al. 2016). Some kinases, for example most receptor tyrosine kinases (Beenstock, et al. 2016) are autophosphorylated in trans, where distinct molecules transfer phosphates to one another. Other kinases autophosphorylate in cis, where a single molecule transfers phosphates onto itself. In the latter case, autophosphorylation of DYRK and GSK3β has been proposed to occur during intermediate stages of protein folding (Lochhead, et al. 2005), by binding to allosteric modulators that reorient the activation loop towards catalytic residues (e.g. p38α, (DeNicola, et al. 2013) or by dimerization driving a conformational change (e.g. RAF, (Rajakulendran, et al. 2009)).

In contrast, for many kinases, activation requires phosphorylation of the activation loop in trans by a distinct upstream kinase. For example, many mitogen activated protein kinases (MAPKs) require phosphorylation by upstream MAPK kinases (MAPKKs) (Chang and Karin 2001). The MAP kinases ERK1 and ERK2 exhibit very low autophosphorylation activity as purified proteins *in vitro* (Levin-Salomon, et al. 2008; Beenstock, et al. 2014), and they manifest high activity *in vivo* only when they are phosphorylated by the upstream MAP kinase kinase, MEK. ERK1 and ERK2 mutants have been described that exhibit much higher autophosphorylation activity (Emrick, et al. 2006), and one mutant is reported to be able to transform the kinase to an oncoprotein (Smorodinsky-Atias, et al. 2016) highlighting the importance of stringent control of ERK phosphorylation.

In earlier work (Howard, et al. 2014), we used ancestral resurrection methods to study the evolution of the specificity of the CMGC group of kinases, which include the cyclin dependent kinase (CDK) and MAPK families. CDK and MAPK are closely related subfamilies within the CMGC group and regulate a wide range of cellular processes. CDKs are major regulators of cell division (Morgan 2007) and transcription (Fisher 2005). MAPKs are crucial for cell proliferation, differentiation, and stress responses. As part of this study, we observed that a deep ancestor of this group had relatively high intrinsic activity even in the absence of any activating kinase of or allosteric activators. This finding was surprising given the tight control of MAP kinases by upstream activators, and the absolute requirement of CDKs on both cyclin binding and upstream activating kinases. Therefore, to study how suppression of autophosphorylation and thus strong dependence on upstream activating kinases evolved, we decided to investigate the regulatory evolution of the CMGC group, ultimately focusing on the MAP kinases ERK1 and ERK2.

Throughout the evolutionary history of MAP kinases, gene duplications and diversifications resulted in multiple paralogs with diverse regulation (Howard, et al. 2014). For example, within the p38 subfamilies, p38 β has a much higher intrinsic catalytic activity than p38α (Beenstock, et al. 2014) and within the ERK family, ERK7 is constitutively active(Abe, et al. 1999; Abe, et al. 2001), and the core kinase domain of ERK5 has a much higher autophosphorylation rate, and thus higher intrinsic activity, than ERK1 and ERK2, which are almost completely inactive in the absence of the activating kinase, MEK (Buschbeck and Ullrich 2005).

Through biochemical analysis of resurrected kinases, we found that the transition to dependence on MEK (a MAPK activating kinase) occurred between the common ancestors of ERK1, 2 and 5 and that of ERK1 and 2. Through a targeted mutagenesis approach we identified the key evolutionary mutations that mediated this transition. Our studies uncovered a role for the gatekeeper residue and the length of the linker loop between the β3 strand and helix αC in this regulatory evolution. Introduction of these evolutionary mutations into modern ERK1 and ERK2 led to full autoactivation in the absence of an upstream activating kinase. Thus, we believe we have identified important steps in the regulatory evolution of the ERK MAPK family.

## Results

### Reconstruction of the lineage leading to ERK kinases

To understand how kinase activity evolved, we resurrected ancestors of the kinases leading to ERK1 and ERK2. Briefly, we collected a set of sequences for each of the major kinase families of the CMGC group. The organisms sampled were selected to be well distributed across the tree of life. We then used a maximum likelihood phylogenetic method (Hanson-Smith and Johnson 2016) to infer the sequences of the ancestral kinase AncCDK.MAPK from which all CDK and MAPK descend, and the common ancestors of the MAPK (AncMAPK), Jnk/p38/ERK (AncJPE), ERK (AncERK), ERK1/2/5 (AncERK1-5) and ERK1/2 (AncERK1-2) families (Figure 1A).

**Figure 1:**
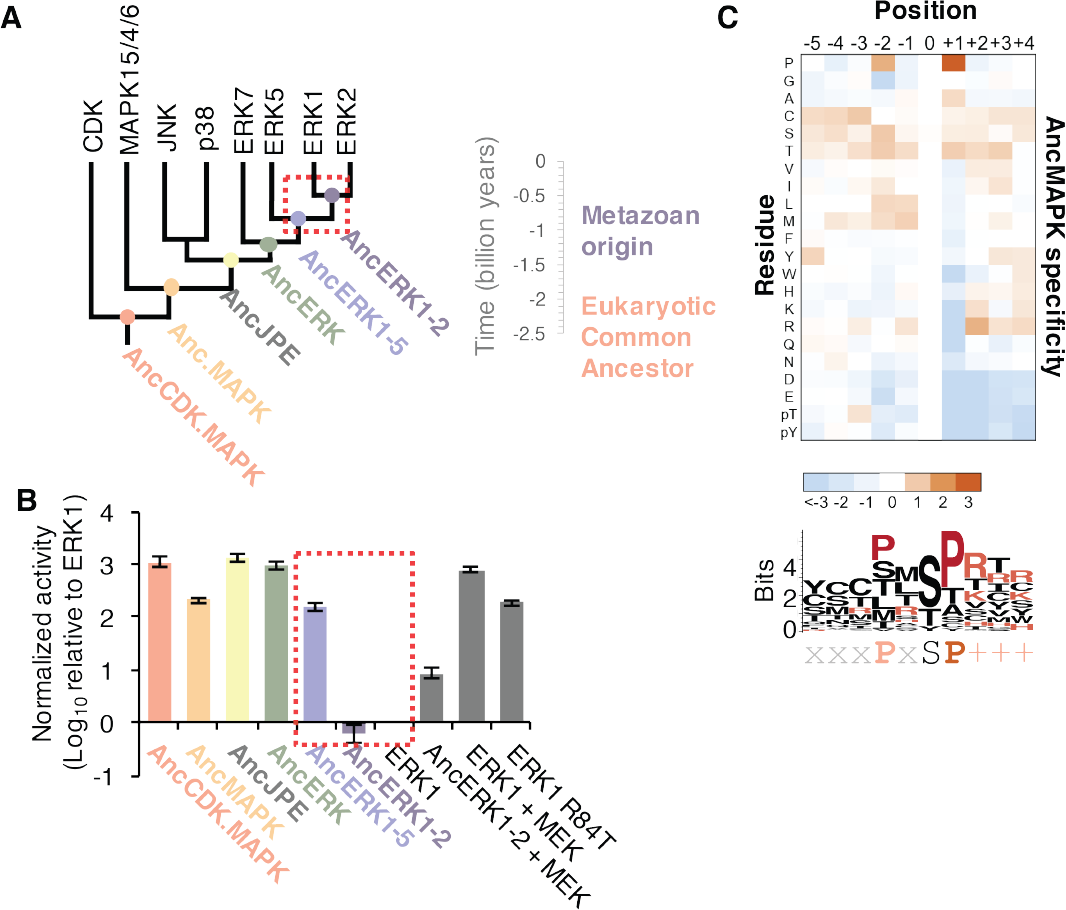
Kinase activity became more tightly regulated at the transition from AncERK to AncERK1-2. (A) Phylogenetic tree for the CDK.MAPK family. The ancestral nodes corresponding to the kinases resurrected in this study are indicated by circles. The red box indicates activity transition from high intrinsic activity to low activity in the absence of activation by MEK. (B) Quantification of the basal activities of ancestral kinases and modern ERK1 with and without the activating kinase, MEK (log_10_ rate of myelin basic protein phosphorylation activity relative to ERK1) the red box indicates the transition in basal activity. (C) Substrate specificity of AncMAPK. Positional scanning peptide libraries were used to profile the specificity of the ancestral kinase AncMAPK. Red indicates positive selection, blue indicates negative selection. The average of two replicates was calculated for each kinase.

### Evolution of substrate specificity in the MAP kinase lineage

We synthesized the coding sequences for these inferred ancestors, and expressed and purified them from *E. coli*. We then used a positional scanning peptide library (Hutti, et al. 2004) to determine the specificity of all ancestors. The specificities of all ancestors were consistent with expectations. As previously reported, the deep ancestor of the CMGC group contained hallmarks of specificity found along distinct branches of the CMGC phylogenetic tree including +1 proline,-2 proline and −3 arginine preference. The new MAPK ancestors generated for the current study lost the −3 arginine specificity at the transition to the ancestor of CDK and MAPK (AncCDK.MAPK, supplemental figure 1A and B). This was most likely due to the mutation of Glu to Ser in the conserved HRDxKPEN motif, previously reported to be a strong determinant of −3 arginine preference (Mok, et al. 2010). All ancestors had a strong preference for proline at the +1 position including the ancestor of all MAP kinases (AncMAPK, Figure 1C), as expected. Together the specificities of ancestors were concordant with expectations providing confidence in our resurrected kinases.

### Regulatory evolution in the MAP kinase lineage

Next, we tested the intrinsic activity of these resurrected kinases. We determined the kinase activity of the ancestors using Myelin Basic Protein (MBP) as a generic substrate. We found that all ancestors prior to and including AncERK1-5 had relatively high intrinsic activity, while the ancestor of ERK1 and ERK2 had low activity, almost comparable to modern ERK1/2 (Figure 1B). Thus, the transition in intrinsic kinase activity occurred between AncERK1-5 and AncERK1-2. Importantly, AncERK1-2 was activated ~10-fold by MEK, indicating that the decrease in intrinsic activity was a regulatory evolution event rather than a general problem with kinase reconstruction leading to loss of activity for trivial reasons. Therefore, we identified a major transition in kinase regulation within the ERK kinase family.

### The higher intrinsic activity of deep ancestors is due to autophosphorylation on the TxY motif

We next wished to determine the mechanism of the regulatory transition between AncERK1-5 and AncERK1-2. Almost all MAPKs have a conserved ‘TxY’motif in their kinase domain, and dual phosphorylation of this motif drives kinase activation. Dual phosphorylation of the threonine and tyrosine residues of ‘TxY’motif in ERK2 is catalyzed by MEK1/2. This dual phosphorylation causes the N- and C-terminal domains in ERK2 to rotate toward one another, thus organizing the catalytic sites (Canagarajah, et al. 1997). This conformation change also promotes substrate recognition in the P+1 site (Figure 2A).

**Figure 2:**
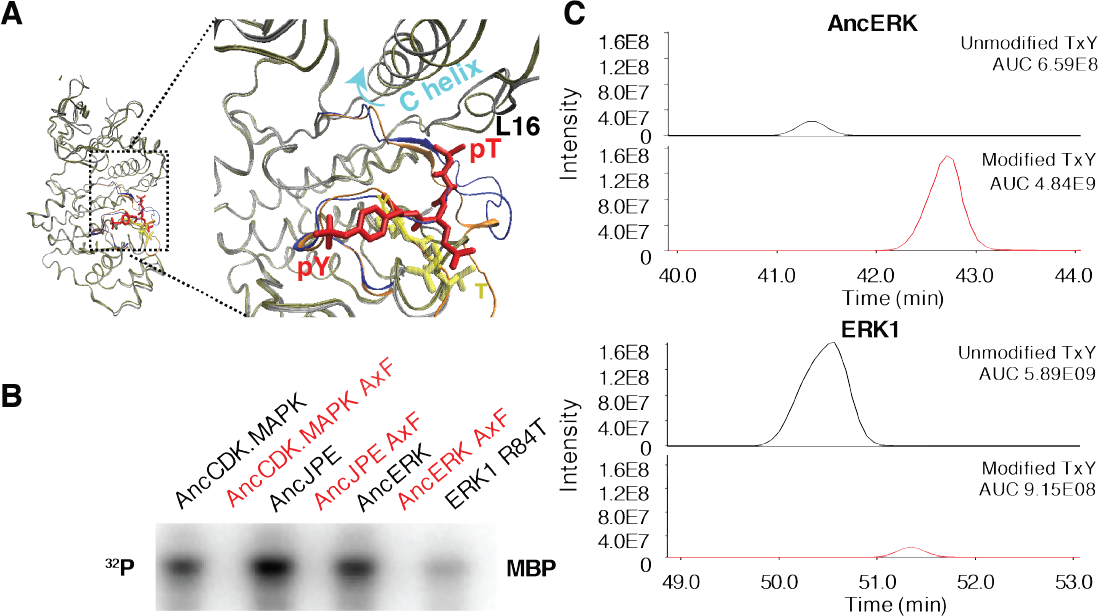
Higher intrinsic activity of ancestors is due to autophosphorylation on the TxY motif. (A) Superposition of ERK2 (silver, Protein Data Bank code 1ERK) and phosphorylated ERK2 (tan, Protein Data Bank code 2ERK). The activation segment (ribbon diagram) and the ‘TEY’ motif (represented as sticks) are colored orange and yellow for ERK2, blue and red for phosphorylated ERK2. Phosphorylation of the TEY motif leads to a conformational change that remodels the activation loop to enable substrate binding, and rotation of the C helix (arrow) leading to interaction between Lys (K52) on the β3 strand and Glu (E69) on the C helix. Part of Loop L16 adopts a helical conformation in phosphorylated ERK2. (B) Kinase activity of wildtype and AxF mutants of ancestors towards MBP. (C) Quantification of phosphorylation level of ‘TEY’ motif in AncERK1 by LC/MS.

Multiple sequence alignment analysis showed that the ‘TxY’motif was conserved among all kinase ancestors (Figure 2-supplement 1A). To test the role of ‘TxY’motif in the evolution of kinase activity, we first mutated the ‘TxY’motif to ‘AxF’, thus removing the phosphoacceptor hydroxyl groups rendering the motif non-phosphorylatable1Q. These mutations greatly reduced the activity of all kinase ancestors (Figure 2B), suggesting the requirement for ‘TxY’phosphorylation was an early evolutionary event. Next we quantified ‘TxY’phosphorylation in AncERK and ERK1 by performing liquid chromatography – mass spectrometry (LC-MS) (Figure 2C; Figure 2-supplement 1B). The phosphorylation level of ‘TxY’motif in AncERK was much higher than that in ERK1, pointing to a high autophosphorylation rate in ancestors. These results indicated that the increased auto-phosphorylation in ‘TxY’in ancestor led to higher intrinsic activity, while a decreased ‘TxY’auto-phosphorylation in modern ERK1 led to lower activity, thus leading to dependence on MEK1/2 for activation.

### Shortening of linker loop and mutation of gatekeeper residue led to tight regulation in the ERK1-2 family

To understand how this regulatory evolution occurred, we next sought to identify the specific mutations underlying the transition in kinase activity that occurred between AncERK1-5 and AncERK1-2. First however, we wished to determine whether the activities of AncERK1-5 and AncERK1-2 were robust to uncertainties in ancestral sequence reconstruction. To address this issue, we synthesized alternative ancestors using a Bayesian sampling approach. The ancestors of alternative kinases behaved similarly to the original ancestors (Figure 3-supplement 1), further suggesting that the transition of kinase activity occurred between AncERK1-5 and AncERK1-2 and that our results are robust to phylogenetic and reconstruction uncertainty (Hanson-Smith, et al. 2010).

With this test, we were confident to compare ancestors in an attempt to identify causal evolutionary mutations. As an initial strategy, we constructed a series of chimeras between AncERK1-5 and AncERK1-2. First, we divided the ancestor kinases into 3 parts (N terminus, middle, and C terminus, Figure 3-supplement 2A and 2B), and shuffled these parts (Figure 3-supplement 2B, chimera 1 − 3). This initial assay indicated that changes in the N-terminus were responsible for most of the difference in the activity of these ancestors (Figure 3-supplement 2D). Next, we narrowed the fragment of AncERK1-5 using two additional chimeras (Figure 3-supplement 2B and 2D, chimera 4 and 5). A key advantage of ancestral resurrection is that the number of amino acid transitions between ancestors is typically far lower that the number of differences between extant proteins (Thornton 2004). Therefore, after our chimera analysis there were only 2 possible changes to assess within the causal fragments, an insertion in the β3-αC loop (44R) and a transition from glutamine to methionine at the gatekeeper residue at the back of the ATP-binding pocket (Q89M) (Figure 3B). Indeed, reverting these amino acids to the ancestral state in AncERK1-2 was sufficient for high basal activity in this kinase (Figure 3A). Conversely, introducing these two mutations into AncERK1-5, suppressed basal activity to the levels of AncERK1-2 (Figure 3-supplement 2C and E). From the Erk2 structure (PDB: 2ERK), the 44R insertion was mapped to the loop between the β3 strand and helix αC, and the Q89M mutation was localized at the gatekeeper position (Figure 3B). Together, these results suggested that the Q89M and 44R insertion were key mutations in the evolutionary transition from high to low intrinsic activity.

**Figure 3:**
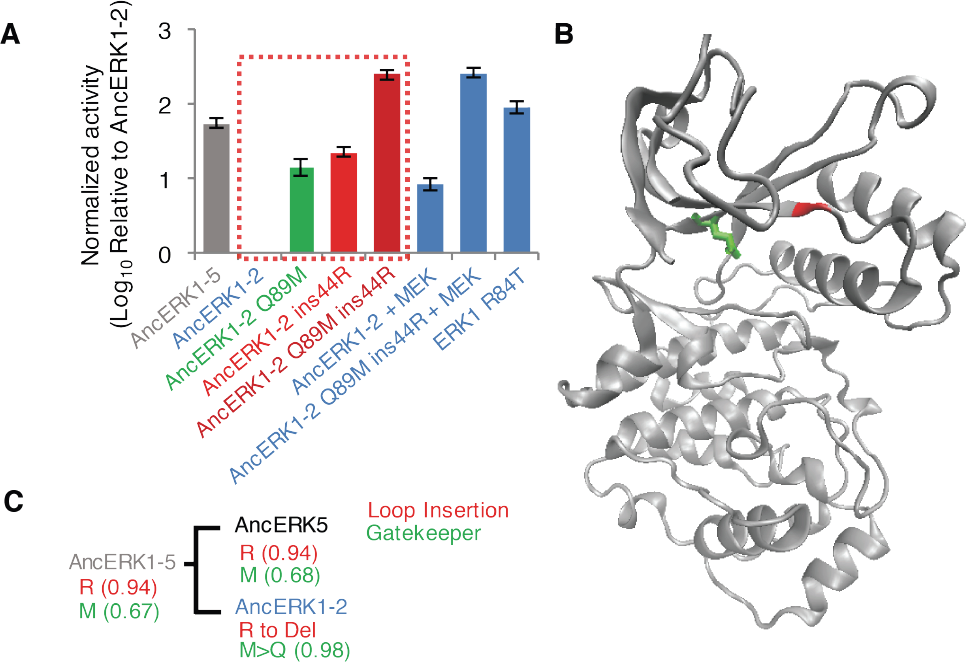
Shortening of the linker loop and mutation of the gatekeeper residue led to tight regulation in the ERK1-2 family. (A) Kinase assays comparing the activity of two evolutionary mutants to the activity of AncERK1-2 before and after preincubation with active MEK kinase (log_10_ rate MBP phosphorylation). (B) Positions of the two evolutionary mutations modeled on the ERK2 structure (PDB 1ERK). The gatekeeper residue, Q122, is shown in green, and the location of the 74N insertion is shown in red. (C) Phylogenetic tree indicating the state of the β3-αC insertion and identity of gatekeeper residues during evolution of the AncERK1-5 group. Numbers in the brackets indicate the posterior probability for ancestral reconstructions.

### Reversing evolutionary mutations leads to activation of modern ERK1

The two mutations that led to low basal activity of ERK1/2 are relatively new innovations. All the deep ancestors had this insert in the linker loop and hydrophobic residue at the gatekeeper (Figure 3C; Figure 3-supplement 3 and 4) and no extant MAPK has this combination of residues. Therefore, we wondered whether mutating these residues back to an ancestral state might be sufficient to increase modern ERK1/2 activity. First, we introduced the insertion into the loop between the β3 strand and αC helix in ERK1. The insertion increased ERK1 activity more than 30-fold (Figure 4A; Figure 4-supplement 1). We also found that the Q122M gatekeeper mutant led to ERK1 activation (figure 4A) and phosphorylation of the ‘TEY’ motif was enhanced (Figure 4B). Mutation in the gatekeeper residue has been reported to activate ERK2 in previous studies (Emrick, et al. 2006), but this mutation was to ALA or GLY, therefore it is interesting that both small residues and the evolutionary mutation to a bulky polar residue both activate ERK1. Interestingly, when we combined the 74 loop insertion with the Q122M gatekeeper mutation, the activity of this double mutant was much higher than either single mutant (Figure 4A). Indeed, the combination of these two mutations raised ERK1 activity by more than 2 orders of magnitude, almost to the levels of ERK1 after activation with MEK. Therefore, reversing just these two evolutionary changes in modern ERK1 reverts the kinase back to an ancestral regulatory state with very little dependence on its upstream regulatory kinase.

**Figure 4:**
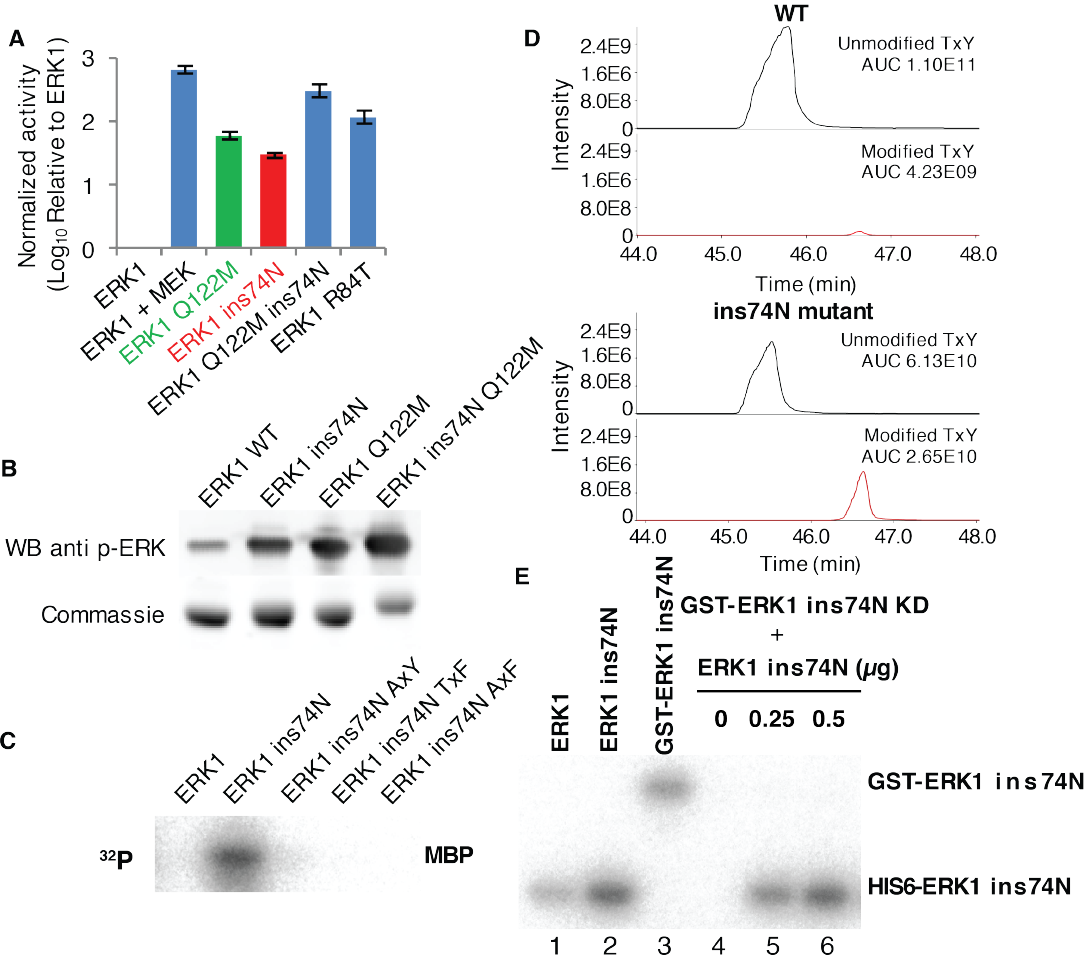
Reversing evolutionary mutations leads to activation of modern ERK1 through cis-autophosphorylation of the TEY motif. (A) Kinase assays measuring activity of modern ERK1 and evolutionary mutants (log_10_ rate of MBP phosphorylation relative to ERK1). (B) Western blots indicating degree of phosphorylation of the residues in the kinase activation loop (top), Coomassie brilliant blue loading control (bottom). (C) Activity of ERK-ins74N mutant and its ‘TEY’ mutants towards MBP. (D) Quantification of the phosphorylation level of the ‘TEY’ motif in ERK1 wildtype and ins74N mutant by LC/MS. (E) Autophosphorylation assay. 0.5μg purified protein of Erk kinase and its mutants were incubated with kinase assay buffer without substrate. The ERK1 kinase and the ins74N mutant were tagged with either His6 (lanes 1 and 2) or GST (lane 3, the GST tag was used to alter the mobility of kinase alleles allowing them to be distinguished on an SDS-PAGE gel). GST-ERK1 ins74N kinase dead K71R mutant (ins74N KD) was incubated without HIS6-ERK1-ins74N (lane 4), and with 0.25μg or 0.5μg HIS6-His6ERK1 ins74N (lanes 5 and 6).

### The length of the β3-αC loop determines activity

Interestingly, in addition to arginine, many other amino acid insertions also increased ERK1 activity (Figure 4-supplement 1), suggesting that this effect was not arginine specific. A repeat of our reconstruction showed that arginine was not a unique insertion at position 44. Asparagine was the next most likely alternative ancestral amino acid at this position. Therefore, we tested an ERK1-74N insertion mutant (ins74N) and determined this mutant to be just as active as the ERK1-74R allele. Thus, we hypothesized that loop-length rather than precise amino acid identity might explain the increased kinase activity. To test this hypothesis, we inserted and deleted residues along this loop. Deletions and insertions from K72 to P75 greatly affected mutant activity. However, length changes after P75, had a minor effect (Figure 4-supplement 2A and 2B). This result confirmed the importance of the loop length in regulating activity and further delimited the loop boundary. Together, these results suggested that the length of the loop between K72 to P75 is an important determinant of basal ERK1 activity.

### Increased activity in modern ERK1 is due to cis autophosphorylation on the TxY motif

Because the ‘TxY’motif was highly phosphorylated in ancestral kinases, we hypothesized that increased ‘TxY’phosphorylation would likely also be a mechanism of mutant ERK1 activation. Consistent with this idea, mutations in this motif to ‘AxY’, ‘TxF’ and ‘AxF’ all abolished the increased activity caused by the insert mutation (Figure 4C). Furthermore, Western blots using an antibody specific for phosphorylated ERK (Figure 4B) and LC-MS quantification (figure 4D; Figure 4-supplement 3B) showed elevated ‘TxY’phosphorylation in mutant kinase. We mostly detected monophosphorylated peptide (Figure 4-supplement 3B), indicating that ‘TxY’phosphorylation is not a processive reaction.

The *E. coli* bacterium that we use to express ERK1 and ancestral kinases lacks endogenous kinases capable of phosphorylating the ‘TxY’motif. Therefore, the increased phosphorylation of ‘TEY’ motif is likely caused by kinase autophosphorylation (Howard, et al. 2014). Indeed, experiments incubating kinase with ATP-y-^32^P in the absence of substrate showed that autophosphorylation rates were indeed increased in the ERK1ins74N mutant (Figure 4-supplement 3A). Together, these results suggested that the insertion caused elevated phosphorylation level of the ‘TxY’motif and increased activity. To test whether this autophosphorylation occurred in cis or in trans, we employed a catalytically inactivated mutant, ERK1-ins74N-K52R, fused to GST to increase the molecular weight of this protein allowing us to distinguish it from catalytically active ERK1 on an SDS-PAGE gel. We then mixed this kinase dead allele with the activated ERK1-ins74N allele. There was no detectable phosphorylation of the kinase-dead GST-ERK1-ins74N-K52R, even though the catalytically active allele clearly continued to undergo autophosphorylation (Figure 4E). This result implies that autophosphorylation occurs through an intramolecular mechanism in cis.

### The catalytic residues are organized in a more active conformation in the insert mutant

We wondered about the molecular mechanism through which the two evolutionary mutations affected kinase activity. We reasoned that since autophosphorylation was increased through an intramolecular mechanism, the activation loop should be more readily phosphorylated by the catalytic residues. There are several critical residues for catalysis: K52, D147 and D165 in the ERK2 numbering (Roskoski 2012). We supposed there might be two possibilities to increase the frequency of ‘TEY’ motif phosphorylation. One was that the activation loop might be more flexible and thus frequently proximal to the catalytic residues. The other was that the catalytic residues might obtain more productive conformation in the mutant, for example the αC helix would rotate to the “in” position, thus better orientating the catalytic aspartate within the conserved DFG motif (Roskoski 2012). To distinguish these two possibilities, we performed a series of molecular dynamic (MD) simulations. Since our kinase assay experiment showed that the loop insertion mutation also increased the activities of both ERK1 and ERK2 to a similar extent (Figure 5-supplement 1A), we could explore the effects of the β3-αC loop insertion mutant in both ERK1 (74N_ins_) and ERK2 (55N_ins_). This afforded the opportunity to inspect the unphosphorylated state, only available for ERK2, as the monophosphorylated starting state, which is only available for ERK1. In each simulation, we assessed both the overall flexibility of the kinase fold and the relative distances between the key catalytic residues in the kinase active site.

For non-phosphorylated ERK2 (PDB ID: 1ERK) (Canagarajah, et al. 1997), root mean square fluctuation (RMSF) calculations that measure the amplitude of residue atom motions during MD simulations did not show significant differences between the overall flexibility of wild type and mutant ERK2 (Figure 5A and 5B). However, we did find changes in the orientation of the β3-strand relative to the αC-helix due to the mutation. The salt bridge between the β3-lysine and αC-glutamate promotes kinase activity and this requires the “αC-in” conformation (Roskoski 2012), thus the distances between the residues β3-K52 (K71 in ERK1) and αC-E69 (E88 in ERK1) are informative for the expected catalytic rate of the active site. The K52-E69 distance in the insertion mutant was decreased relative to WT (Figure 5C and Figure 5-supplement 1B), suggesting that the catalytic salt bridge K52-E69 was more easily formed in mutant. This result is consistent with the idea that the catalytic site of the unphosphorylated enzyme was more efficient and thus autophosphorylation of the activation loop would occur more readily.

**Figure 5:**
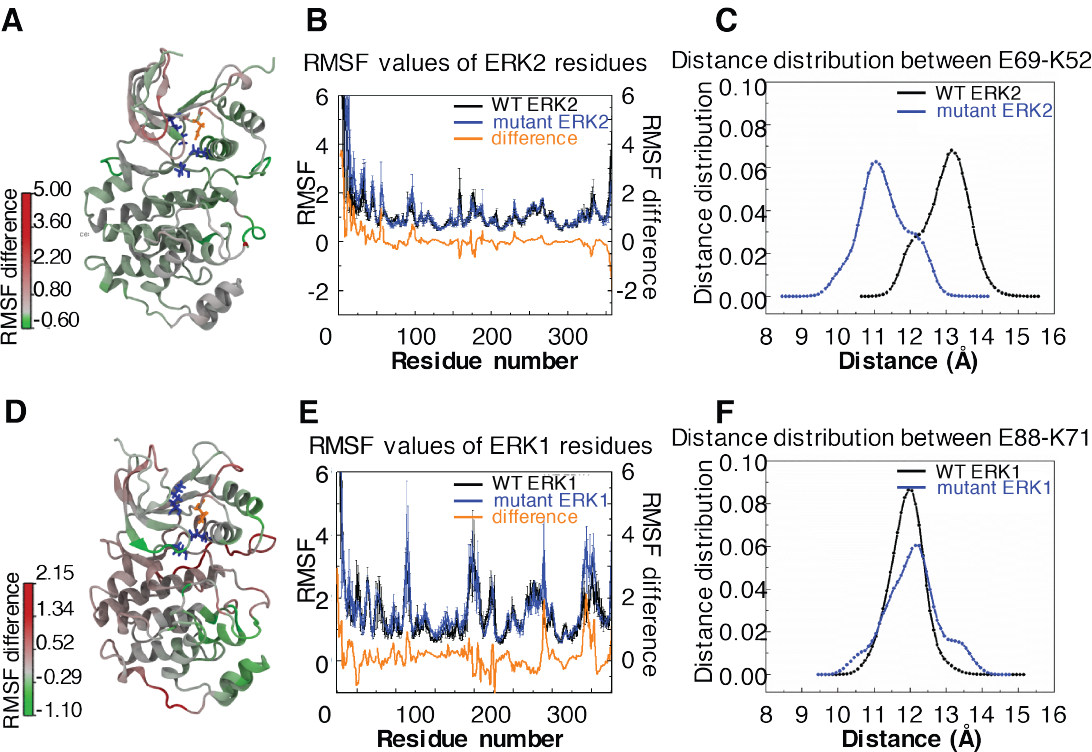
Molecular Dynamics simulations show rearrangement of the active site in unphosphorylated ERK2 and increased mobility of the L16 loop in monophosphorylated ERK1. (A, D) RMSF differences between WT and mutant ERK projected onto on crystal structures 1ERK (Erk2, A) and 2ZOQ (Erk1, D). The catalytic residue lysine on the β3 strand is indicated in orange. The Glutamate on the C helix, aspartate in the HRD motif, and aspartate of DFG are indicated in blue. (B, E) RMSF values for WT (black) and insertion mutant kinases (blue), and the difference between WT and mutant (orange) were plotted for each residue. (C, F) Distributions of distances between the C helix glutamate and β3 strand lysine in WT (black) and mutant (blue) kinases.

### The insertion mutant may facilitate domain closure in the monophosphorylated kinase

We also performed MD simulations of the monophosphorylated ERK1 proteins to identify the effects of the ins74N mutation on WT ERK1 (PDB ID: 2ZOQ) dynamics (Figure 5D). In this case, the distance between the residues-K71 and E88- (equivalent to the K52 and E69 in ERK2) showed no significant difference between WT and insertion mutant kinase (Figure 5F and Figure 5-supplement 1C). However, the L16 loop was significantly more flexible in the insertion mutant (Figure 5D and 5E). Previous MD simulations have implicated an increased frequency of domain closure as part of the mechanism of increased activity in the gatekeeper mutant kinase (Barr, et al. 2011). Refolding of the L16 loop is involved in domain closure, and so this result suggests a mechanism for the synergy between the gatekeeper and loop insertion mutations.

### Reversing evolutionary regulatory mutations leads to MEK independent ERK activity *in vivo*

Next we sought to determine whether the evolutionary insertion mutation led to higher ERK1 activity *in vivo*. We generated lentivirus encoding ERK1-WT or ERK1-ins74N and transduced NIH 3T3 cells with these two alleles. We used an ERK-KTR reporter (Regot, et al. 2014), as our reporter for ERK kinase activity (Figure 6A). This reporter contains an ERK docking site and phosphoacceptor sites within nuclear localization and export sites. The reporter is designed such that it is localized in the cytosol when phosphorylated by ERK, and is nuclear in the basal state with low ERK activity. Before imaging, cells were pretreated with the MEK inhibitor Trametinib to prevent activation by endogenous mechanisms. We then performed time lapse imaging of cells to observe the localization of the ERK1-KTR reporter. As expected, we found that the ERK-KTR reporter was mainly localized in nucleus in ERK1-WT cells, and only a small percentage of cells indicated any ERK activity (Figure 6B and 6C), suggesting ERK activity was low after inhibition of MEK by Trametinib. In cells expressing ERK1-ins74N, however, there was significantly more cytoplasm-localized-KTR. This result suggests that the evolutionary mutation led to ERK1 activity independent of MEK *in vivo*.

**Figure 6:**
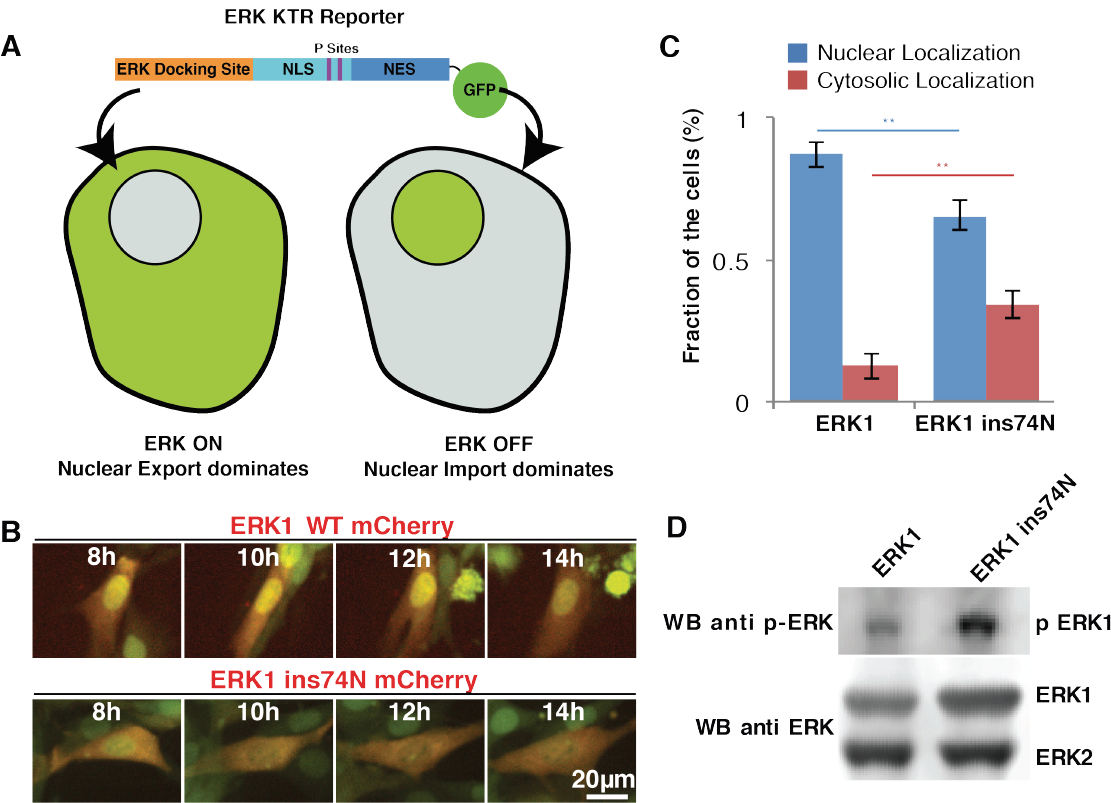
Converting ERK regulatory mutations to the ancestral state leads to MEK independent activity *in vivo*. (A) Kinase Translocation Reporter (KTR): phosphorylation of the reporter by ERK1/2 leads to inactivation of a nuclear localization signal (NLS) and activation of a nuclear export signal (NES), thus leading to nuclear export. Conversely, in cells with low ERK1/2 activity, the NLS is active and NES inactive, therefore the reporter is localized to the nucleus. (B) 3T3 cells expressing ERK1 (indicated by coexpression of DsRED with an IRES, within the ERK1 construct) and the ERK-KTR (green) were treated with 1μM MEK inhibitor Trametinib), and grown without FBS. (C) Statistical analysis of ERK-KTR localization after 12-14 h Trametinib treatment and FBS deprivation. The fraction of cells in which the majority of ERK-KTR localizes to the nucleus or cytosol is plotted. Statistical comparisons are by Student’ s t-test, **^**^**p<0.01. Experiments were repeated three times (N~50 cells per line for each replicate). (D). Western blot to assess phosphorylation of the ERK TEY motif after 14 h growth with Trametinib (1μM) and without FBS (top) and total ERK loading control (bottom).

We also analyzed the phosphorylation level of the two ERK variants *in vivo*, by western blot using an antibody specific to phosphorylated ERK. NIH 3T3 cells transduced with both wildtype and mutant ERK were treated with Trametinib. After treatment, proteins were extracted for western blot. The western blot result showed that the phosphorylation level of ‘TEY’ motif in wild type ERK1 was very low after treatment with Trametinib, while the cells expressing ERK1-ins74N exhibited robust phosphorylation (Figure 6D). These results are consistent with the KTR reporter results, and together show that reversal of the evolutionary insertion mutation led to a decreased dependence on the upstream activating kinase MEK for ERK activity *in vivo*.

## Discussion

### Regulatory evolution in the MAP kinase group

Over several decades, biochemical, genetic and structural studies have provided deep insights into the regulation of kinases. However, the mechanisms by which this diverse array of regulatory mechanisms evolved from a common ancestral kinase remains poorly understood. In this study, we analyzed the activity and autophosphorylation level of the ERK kinases that lie within the CMCG group. During evolution, the intrinsic ability of these kinases to autophosphorylate at their activation loop sharply decreased at the split between the ERK5 and the ERK1-2 kinases. While the intrinsic activity of the ERK5 family remained relatively high, the ERK1-2 kinase became strongly dependent on the upstream activating kinase MEK, part of the Ras-Raf-MEK-ERK pathway that is frequently activated in cancer. In the current study, we took an ancestral resurrection approach to elucidate the likely evolutionary changes that account for this regulatory diversification. We identified two mutations that together account for most of the difference between modern ERK1-2 and ERK5 and reversal of these evolutionary mutations in modern ERK1 led to higher kinase activity and autophosphorylation (Figure 4).

### Mutations in the loop are cancer alleles in other kinase families

It has recently been discovered that changes in the β3-αC loop length of BRAF, EGFR, and HER2, none of which are in the CMGC group of kinases, also led to activation of these kinases (Foster, et al. 2016). In these cases, the activating mutations shorten the loop rather than lengthen it. This probably relates to pulling in the α helix favoring the αC in conformation in these cases. It is interesting that ERK has the opposite behavior. What is generally clear is that the β3-αC loop is a crucial determinant of the kinase activation mechanism.

In previous studies on the evolution of specificity within the CMGC kinase group, we found that an ancestral DFG+1 mutation was also sampled in cancer mutations throughout the kinome (Howard, et al. 2014), therefore it is possible that our evolutionary studies within the CMGC group of kinases may be more generally reflected throughout the kinome. The decreasing confidence in reconstructions with evolutionary distance makes it impossible to resurrect an ur-kinase, but features of this ultimate ancestor are certain to be retained across every kinase group. Thus, we speculate that β3-αC loop length was of critical importance since deep evolutionary history.

Why, then, is ERK not a major oncogene? The upstream kinases BRAF and MEK are frequently activated. Indeed, the Ras/RAF/MEK/ERK pathway is one of the most important signaling axes in oncogenesis. One possibility is that the upstream kinases have multiple targets in addition to ERK, and indeed this is certainly the case for Ras. Another intriguing possibility is that the dynamical properties of ERK signaling are crucial for its role in oncogenesis. ERK is not simply an ON/OFF switch, but rather oscillates or adapts in various contexts. It will be interesting to leverage our mutants and investigate more recent ancestral states that may have distinct dynamical properties to investigate this latter possibility.

### Evolutionary mutations synergize

We found that the evolutionary changes that we determined to drive this increased stringency of regulation synergize. One change, a polar substitution in the “gatekeeper” residue of the active site has been previously described as an activating mutation (Emrick, et al. 2006; Azam, et al. 2008). Evolution at the gatekeeper after AncERK1-5 proceeded along two divergent paths. The permissive methionine residue was retained along the branch that led to the modern ERK5 family in which the acquisition of a large C-terminal domain is thought to be an alternative source of regulation (Buschbeck and Ullrich 2005). In ERK1-2 the gatekeeper was shifted to glutamine, a more restrictive state for autoactivation.

The second evolutionary change that we identified, a change in loop length between the β3 strand and αC helix, has not been observed before. The deeper CMGC ancestors and modern ERK5 all had one extra amino acid in this loop (Figure 3C and Figure 3-supplement 4). The reversal of the shortening of this loop in AncERK1-2 and in modern ERK1 drastically increased kinase activity (Figure 3A and Figure 4).

When combined, these two evolutionary changes synergize to give greater than two orders of magnitude increase in the activity of modern ERK1 and ERK2. Thus, these changes provide an evolutionary pathway of increasing regulatory stringency that may have allowed the system to adapt without any major fitness barriers.

### Structural dynamics and control of kinase activity

We focused our molecular dynamic simulations on the novel β3-αC loop insertion mutation as similar experiments had been performed previously for the gatekeeper mutation (Barr, et al. 2011). We started from either unphosphorylated or monophosphorylated ERK1/2 structures as starting points for these simulations. In the unphosphorylated state, we didn’ t observe any major difference in the flexibility of the kinase domain. However, the distance between K54 and E71 catalytic residues was shorter in the insertion mutant than in wild type (Figure 5C and Figure 5-supplement 1B). It is known that kinase activity requires the formation of this salt-bridge (Roskoski 2012), therefore though encounters between the activation lip and the active site are rare, they would more frequently be productive due to this increase in k_cat_.

MD simulations using monophosphorylated ERK1 as a starting point showed a single significant difference in flexibility at the L16 loop. The reorganization of this loop is important in the domain closure of kinases, which is an intrinsic part of the conformational shift to an active state. This observation is consistent with earlier MD simulations inspecting gatekeeper mutations which also show an increase in domain closure (Barr, et al. 2011), and perhaps explains how these two mutations act synergistically.

### Robust evolution in the CMGC family of kinases

Previous ancestral resurrection studies have highlighted the importance of epistasis in the diversification of function. For example, changes in the specificity of nuclear receptor ligand binding domains required stabilizing mutations (Thornton, et al. 2003) prior to the advent of structure changing mutations, and the acquisition of novel mutations after acquisition of a new function made it difficult to revert to the ancestral state (Bridgham, et al. 2009). In our previous study on the evolution of kinase specificity, we found no requirement for stabilizing mutations – full activity was maintained as specificity changed. In the current study, we find that the reversion of a single or double evolutionary mutation is also well tolerated. Together these observations suggest that kinases are quite tolerant of mutations, which perhaps explains their frequent dysregulation in diseases like cancer. The diversification of specificity and regulation may also be underpinned by this surprising tolerance of a dynamic domain to mutations that drive neofunctionalization. Perhaps this robustness explains the prevalence of kinases as regulatory enzymes in *Eukaryotes*.

## Materials and methods

### Protein Expression and Plasmid Construction

pET28b vectors were used for bacterial expression. Open Reading Frames (ORFs) of the Erk proteins were fused N terminally to the 6X histidine tag for purification. Glutathione S Transferase (GST) was inserted between the 6X His-tag and the Erk ORF in order to increase protein molecular weight. For site-directed and domain-swapping mutagenesis, primers containing the point mutation sites or domain junction sites for chimera protein were designed. The PCR fragments and the vectors were assembled according to the manufacturer’ s instructions using Gibson Assembly Master Mix kit from New England Biolabs (NEB). All the constructs were verified via DNA sequencing.

### Mammalian Cell Culture

NIH3T3 and HEK293T cells were cultured in Dulbecco’ s Modified Eagle’ s Medium (DMEM) supplemented with 10% Fetal Bovine Serum (FBS), penicillin, and streptomycin. The cells were incubated at 37°C and 5% CO2. Transfections of HEK293T cells were performed using TransIT-293 Transfection Reagent (Mirus), according to the manufacturer’ s instructions. Viral supernatants were collected 48–72 h following transfection of 293T cells. After virus infection, NIH3T3 cells were selected with 1-2 μg/ml puromycin. For western blot, cells were first washed 3 times with PBS, then incubated in medium with no serum and treated with 2 μM Trametinib. After 12 hours, cells were lysed by addition of SDS loading buffer containing 20% glycerol, 4% SDS, 0.1 M Tris (pH 6.8), 0.2 M dithiothreitol (DTT), and phenol blue dye, followed by boiling at 100°C for 10 min.

### Protein purification

Protein purification from *E. coli* cells was performed as previously described (Howard, et al. 2014). Briefly, proteins were expressed in Rosetta2 DE3 competent cells by induction with 100 μM IPTG for 18 hr at 16°C. Bacterial culture were collected and centrifugated at 4000rpm for 20 min at 4°C. The cell pellet was resuspended in cold lysis buffer (50mM NaH2PO4, 300mM NaCl, 10 mM imidazole pH7.6, 1 mM PMSF). After sonication, the lysate was centrifuged at 12000rpm for 20 min at 4°C. The supernatant was mixed with magnetic Ni-NTA beads (Biotool) and incubated for 1 hour at 4°C. The bound beads were collected and rinsed 3 × with wash buffer containing (50mM NaH2PO4, 300mM NaCl, 40 mM imidazole pH7.6). The bound proteins were eluted with elution buffer (50mM NaH2PO4, 300mM NaCl, 500 mM imidazole pH7.6). The elution was dialyzed into storage buffer (150mM NaCl, 50mM HEPES pH8, 1mM DTT, 10% glycerol) using Zeba Spin Desalting Columns (Thermo scientific), and flash frozen in aliquots with liquid nitrogen and stored at 80°C.

### Western blot

For recombinant proteins, 200 ng protein were mixed with SDS loading buffer and boiled at 100 °C for 10 min. For mammalian cells, the protein lysates were prepared by removing the medium, washing with PBS, and adding SDS buffer directly to the plate. Cell lysates were collected and boiled for 10 min. The samples were separated on 4-12% Bis-Tris gels (ThermoFisher Scientific) and transferred to a nitrocellulose membrane. The membranes were probed with antibody anti-ERK (SC154, Santa Cruz Biotechnology) or anti-p-ERK (SC7383, Santa Cruz Biotechnology).

### In Vitro Kinase assay

0.05 μg purified recombinant 6x His-tagged Erk protein were used in a total volume of 10 μl. Final reaction conditions were 20 mM Hepes, pH 7.5, 10 mM MgCl_2_, 50 mM NaCl, 0.75 μg/μl myelin basic protein (MBP), 100 μM ATP, and 0.05 μCi of [γ-^32^P]ATP. The kinase reactions proceeded for 30 min at room temperature and were terminated by addition of 6 μl 5x SDS loading buffer (10% SDS, 0.5 M DTT, 50% glycerol, 0.25% Bromophenol blue). All samples were separated on 4-12% Bis-Tris gels (ThermoFisher Scientific). The gel was dried and exposed to phosphor screen. Phosphor screens were analyzed with a Typhoon 9500 scanner (GE) using ImageQuant software (GE).

### Autophosphorylation assay

0.1 μg purified recombinant 6x His-tagged Erk protein was incubated in kinase reaction buffer (20 mM Hepes, pH 7.5, 10 mM MgCl_2_, 50 mM NaCl, 25 μM ATP, and 0.05 μCi of [γ-^32^P]ATP) without the substrate, MBP in a 10μl total volume. The autophosphorylation reactions proceeded for 30 min at room temperature and were terminated by addition of 6 μl 5x SDS loading buffer. All samples were separated on 4-12% Bis-Tris gel (ThermoFisher Scientific). The gel was dried and exposed to phosphor screen. Phosphor screens were analyzed with a Typhoon 9500 scanner.

### Quantification analysis by Liquid Chromatography MS

To extract the protein, a slurry of R2 20 μm Poros beads (Life Technologies Corporation) was added to each sample. The samples shook at 4 °C for 2 hours. The beads were loaded onto equilibrated C18 ziptips (Millipore). Gel pieces were rinsed three times with 0.1% TFA and each rinse was added to its corresponding ziptip followed by microcentrifugation. The extracted poros beads were further washed with 0.5% acetic acid. Peptides were eluted by the addition of 40% acetonitrile in 0.5% acetic acid followed by the addition of 80% acetonitrile in 0.5% acetic acid. The organic solvent was removed using a SpeedVac concentrator and the sample reconstituted in 0.5% acetic acid. 1 μg each sample was analyzed. LC separation online with MS was used with the autosampler of EASY-nLC 1000 (Thermo Scientific). Peptides were gradient eluted from the column directly to QExactive mass spectrometer using a 1 hour gradient (Thermo Scientific). High resolution full MS spectra were acquired with a resolution of 70,000, an AGC target of 1e6, with a maximum ion time of 120 ms, and scan range of 400 to 1500 m/z. Following each full MS twenty data-dependent high resolution HCD MS/MS spectra were acquired. All MS/MS spectra were collected using the following instrument parameters: resolution of 17,500, AGC target of 5e4, maximum ion time of 250 ms, one microscan, 2 m/z isolation window, fixed first mass of 150 m/z, and NCE of 27. MS/MS spectra were searched using a standalone Byonic and Byonic within Proteome Discoverer 2.1.

### Imaging and image analysis

For the construct to measure the phosphorylation of Erk in mammalian cells, KTR ORF was placed under CMV promoter in vector pLentiCMV (Addgene #59150). To create the construct to track Erk expressing cells in KTR experiment, IRES and DsRed was placed after the ORF of Erk1. Cells were seeded on 6 well plates one day before imaging. When the cells were grown to 30% confluency, media was changed to media without FBS and containing Trametinib (final concentration 2 μM) 2 hours prior to imaging. Cells were imaged with a Nikon Eclipse Ti fluorescence microscope. All imaging was carried out at constant conditions of 37°C, 5% CO2, and humidity. The subcellular distribution of the KTR reporter was analyzed by Image J. The cells expressing both Kinase and KTR reporter were used for analysis and the cells that were out of focus and short lived were omitted from analysis.

### MD simulation

In the simulations of Erk1-ins74N and Erk2-ins55N insert mutants, we extracted the structures of Erk1 (PDB ID: 2ZOQ) and Erk2 (PDB ID: 1ERK) from the Protein Data Bank (PDB). We introduced the insertion mutation to both Erk1 and Erk2 using Discovery Studio 4.5 software, (DS) (Biovia, 2015) and then modeled the mutated loops in Erk1-ins74N and Erk2-ins55N structures using Modloop web server (Fiser, et al. 2000). For MD simulations, we used NAMD 2.11 software (Phillips, et al. 2005) with CHARMM22 force field (Mackerell, et al. 2004). First we performed energy minimization of the two initial models. Then we solvated each resulting model in a TIP3P water box with 12Å box padding and added neutralizing ions. We carried out all MD simulations with periodic boundary conditions and using N, P, T ensemble where we kept the temperature constant at 310K using Langevin dynamics with a damping coefficient of 2ps^-1^ and kept the pressure constant at 1atm using The Nose-Hoover Langevin piston method with a 200fs piston period and 100fs decay time. The integration step was 2fs and the cutoff for Van der Waals interactions was 12Å. For computation of long-range electrostatic interactions we used the full particle-mesh Ewald method with 1Å grid spacing and direct space tolerance of 10^−6^. In MD simulations of solvated models, we applied minimization and equilibration for 10,000 steps and 500,000 steps, respectively. We, then, performed 400 ns simulations. We eliminated all rotational and translational motions by aligning the simulation trajectories to the first frame with VMD software 1.9.2 (Humphrey, et al. 1996). We also used VMD for visualization of the simulation trajectories.

### Ancestral Resurrection of the ERK1 and ERK2 lineage

We identified orthologs of human ERK1, ERK2, ERK5 and ERK7 using a protein BLAST (Altschul, et al. 1990) search against the genomes of a diverse set of eukaryotes. The obtained sequences were clustered using CD-HIT (Li and Godzik 2006) at 95% identity to collapse closely related kinases into one sequence. These were then reverse blasted against the human ERKs to identify the ERK that they were most closely related to. For the purposes of reconstruction, they were considered to be the ortholog of this ERK. After some manual screening, the final list of 150 sequences was fed into Phylobot (Hanson-Smith and Johnson 2016) for ancestral resurrection with sequences from similarly collected p38 and JNK orthologs used as an outgroup to root the tree.

## Acknowledgements

We thank Conor Howard for initial experiments and contributions to this project, David Truong for help writing the manuscript; Greg Brittingham and Morgan Delarue for help with imaging and analysis; Laura Hug and Victor Hanson-Smith for advice regarding ancestral resurrection; and David Engelberg for sharing plasmids. We also thank Yu Zhao and the rest of the Holt lab for helpful discussions regarding the project. We also thank our core facilities, particulary Beatrix Uberheide and the staff at the proteomics core at NYU Medical Center. We gratefully acknowledge funding from the William Bowes Fellows program and the Vilcek Foundation (LJH).

---

**Figure 1-figure supplement 1:**
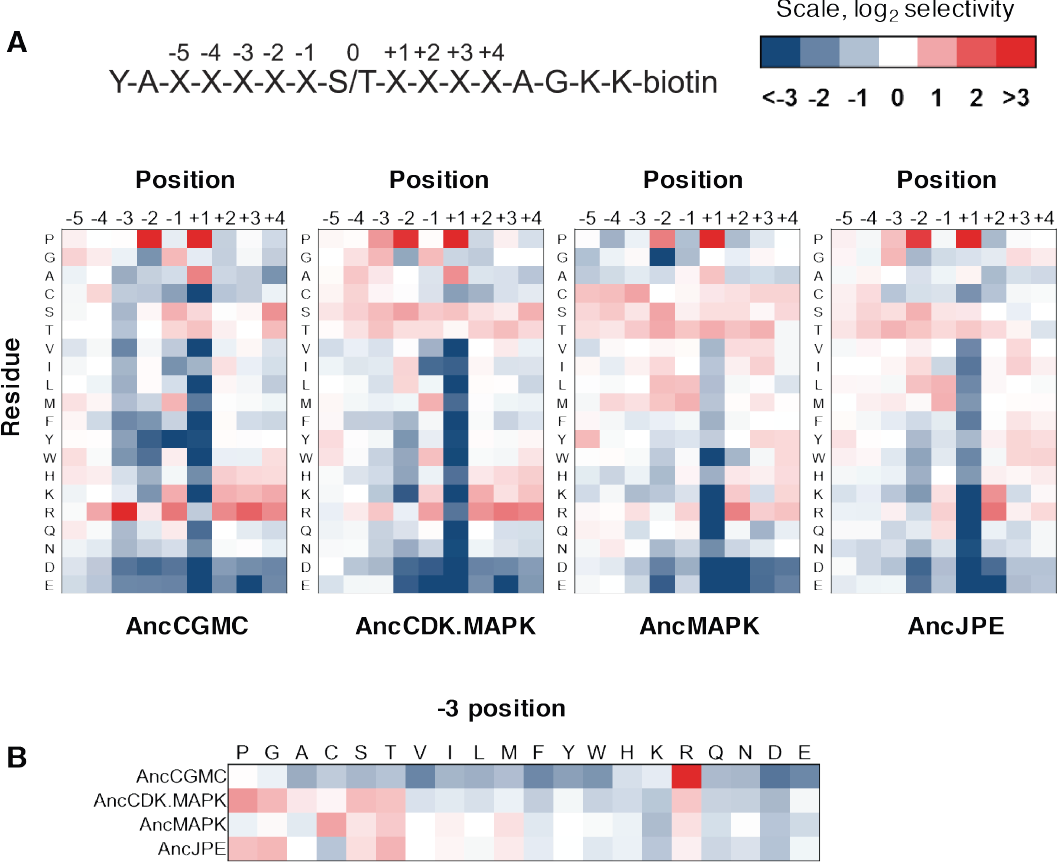
Determination of the primary sequence specificity of ancestral kinases. (A) The specificity of AncCGMC, AncCDK.MAPK, AncMAPK and AncJPE were profiled by scanning peptide libraries. Red indicates positive selection for a given amino acid while blue indicates negative selection. A schematic of the peptide library is shown above. The positions are numbered according to the schematic of the peptide library. (B) A subset of the data above focusing on the evolution of −3 selectivity, with diminishing preference for arginine in more recent kinases.

**Figure 2-figure supplement 1:**
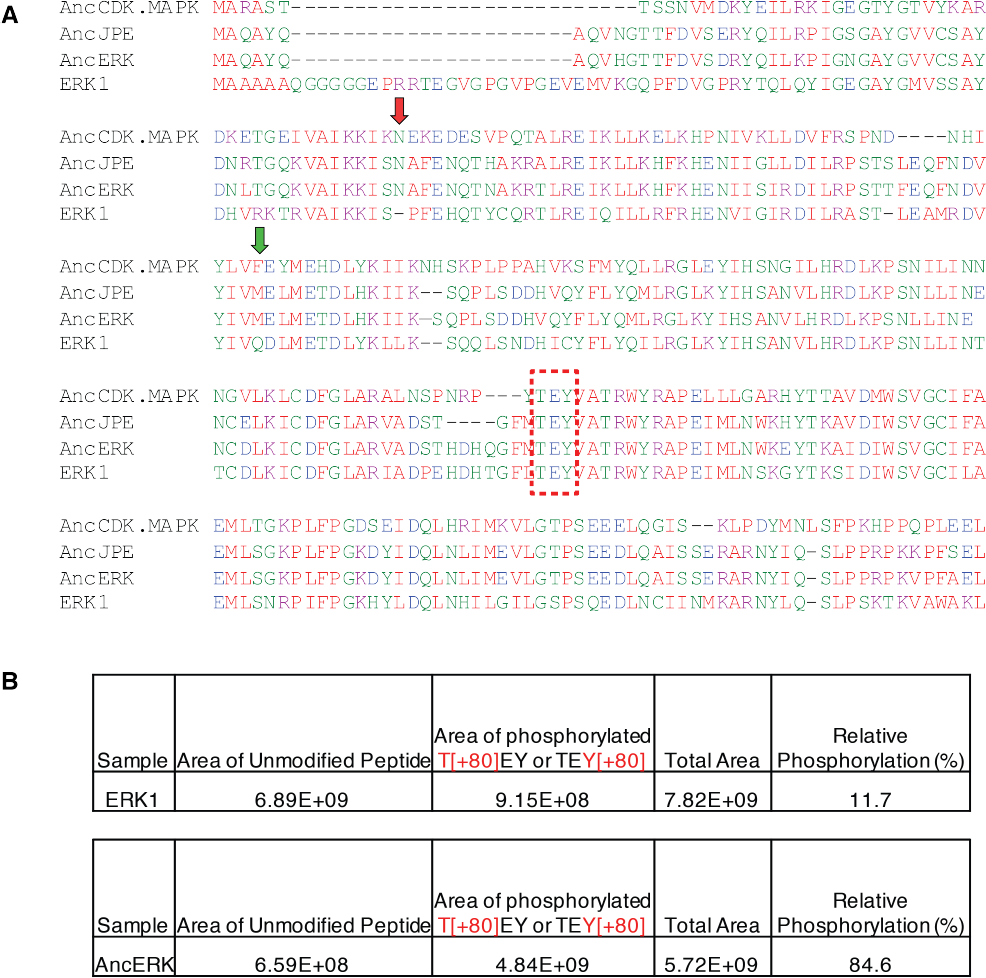
Sequence alignment of ERK1-2 kinase ancestors and modern ERK1 and MS quantification of ‘TEY’ phosphorylation in ERK1 and AncERK. (A) Primary sequences of the core kinase domains of ancestors and modern ERK1 were aligned using MUSCLE. The ‘TEY’ motif is indicated by the red box. The red and green arrows indicate the evolutionary mutations that led to decreased dependence of AncERK on MEK. (B) Quantification of the phosphorylation level of ‘TEY’ motif in AncERK and ERK1 by LC/MS. The area under the peak corresponding to the peptide containing ‘TEY’ in Erk1, and AncERK was quantified. No di-phosphorylated peptide was detected. The areas of phosphorylated threonine and tyrosine are combined in the table.

**Figure 3-figure supplement 1:**
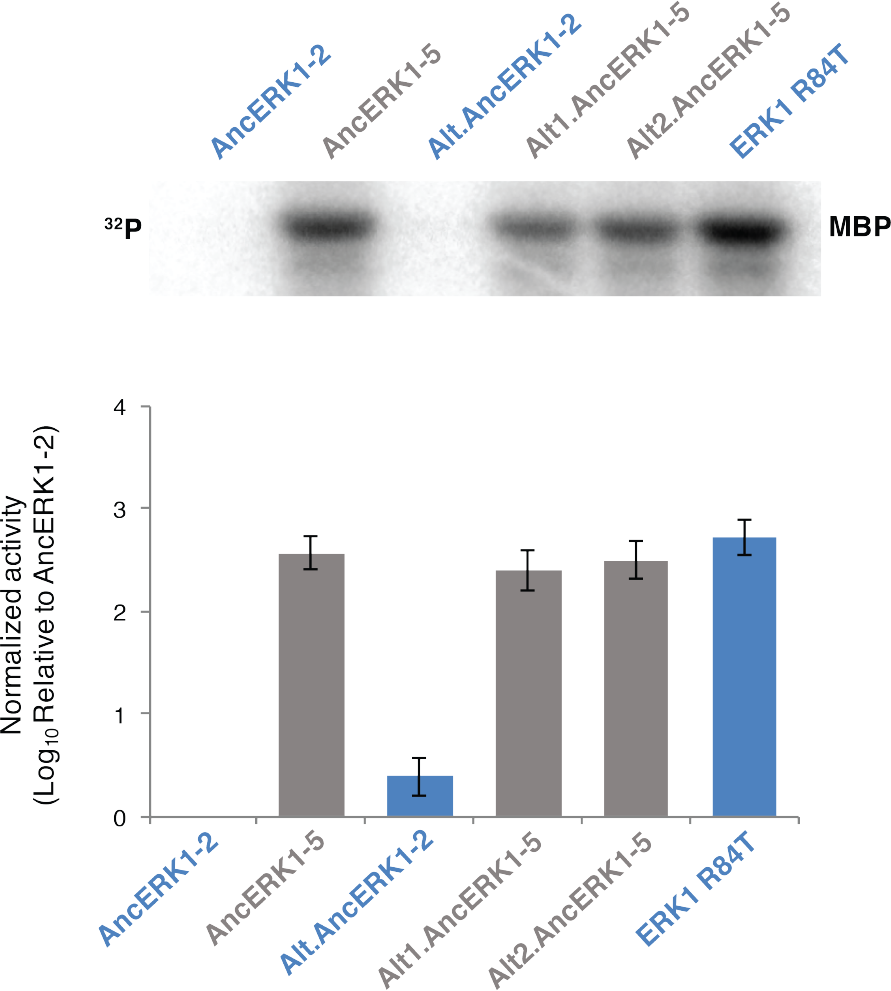
Activities of AncERK-2 and AncERK1-5 are robust to uncertainties in ancestral reconstruction. Kinase assays measuring activity of two alternative ancestors of AncERK1-5 and AncERK1-2, normalized to the activity of AncERK1-2 (log_10_ scale). Plausible alternate states with PP >0.2 were all included in the alternate reconstructions (Alt.AncERK1-2 and Alt.AncERK1-5. As there were two alternate states with PP>0.2 at the residue 34 in AncErk1-5, two alternative ancestors of AncERK1-5 were constructed (Alt1.AncERK1-5 and Alt2.AncERK1-5).

**Figure 3-figure supplement 2:**
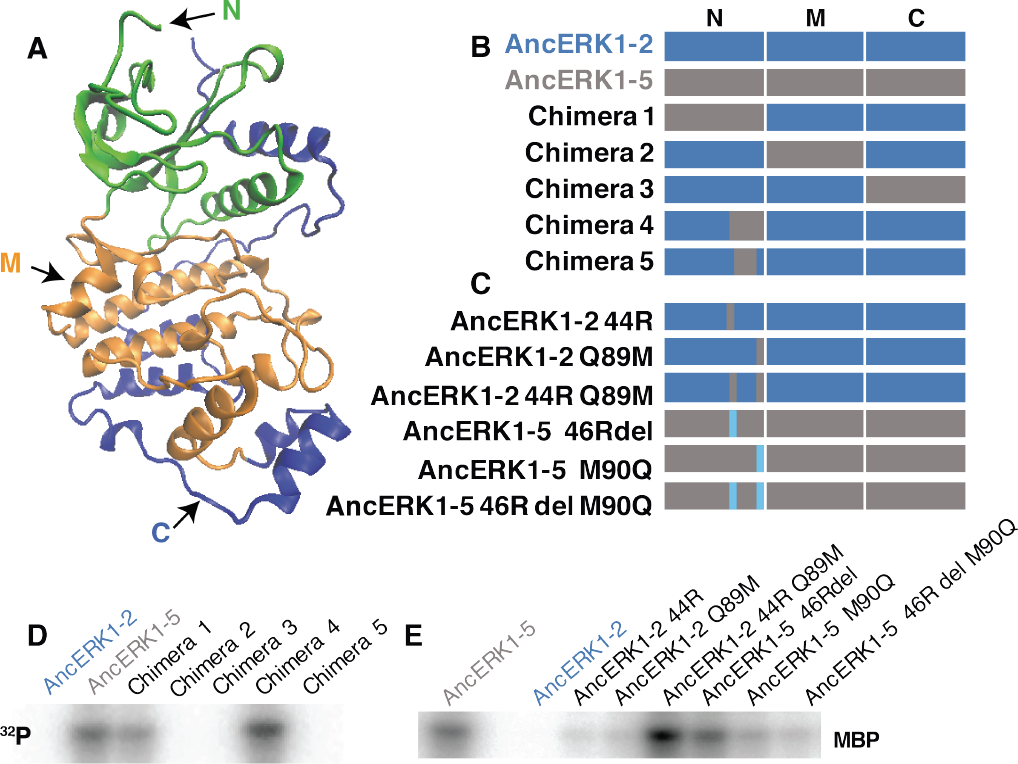
Identification of the evolutionary mutations mediating the decrease in intrinsic activity between AncERK1-5 and AncERK1-2. (A) Kinase domains were divided into N-terminal (green), middle (orange) and C-terminal sections (blue). (B and C) Summary of the chimeric constructs and mutations used to map the mutations that caused a change in the intrinsic activity of ancestors. Blue blocks derive from AncERK1-2, gray blocks from AncERK1-5. (D and E) Kinase assays to determine the intrinsic activity of the chimeras and mutants towards MBP.

**Figure 3-figure supplement 3:**
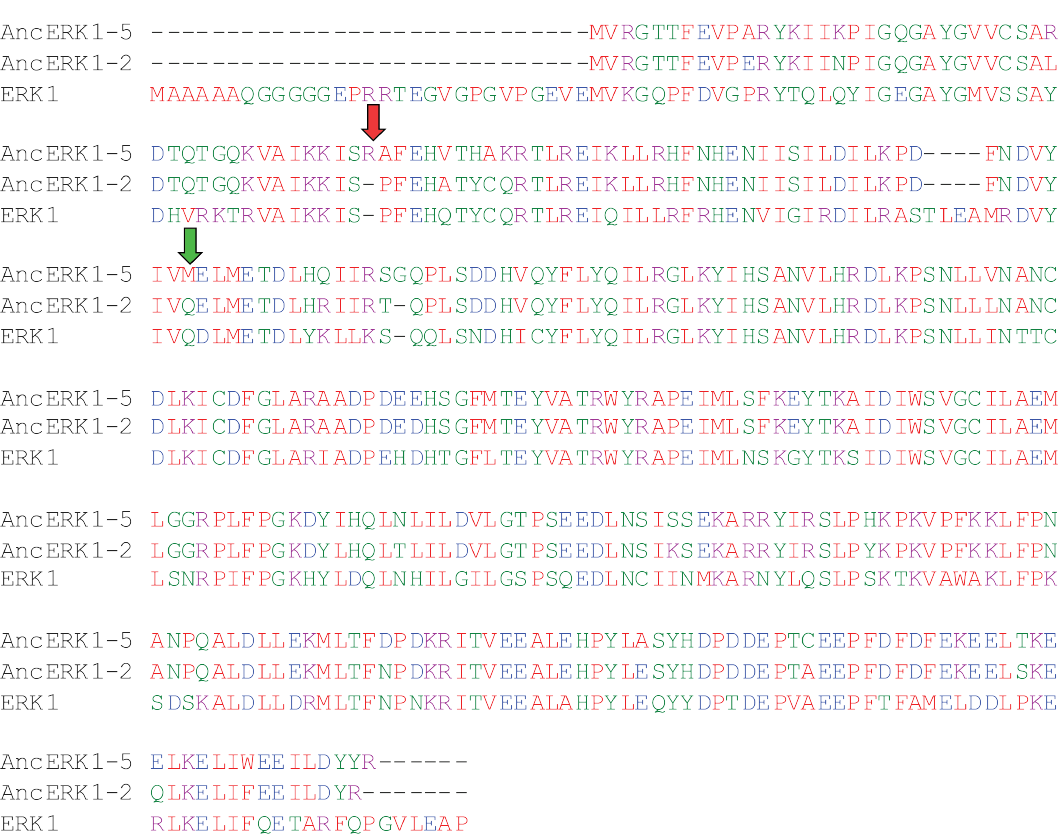
Alignments showing the position of the mutations that drove the transition in activity that occurred between AncERK1-5 and AncERK1-2. The insertion in the β3-αC loop is indicated by a red arrow, while the mutation at the gatekeeper residue is indicated by a green arrow.

**Figure 3-figure supplement 4:**
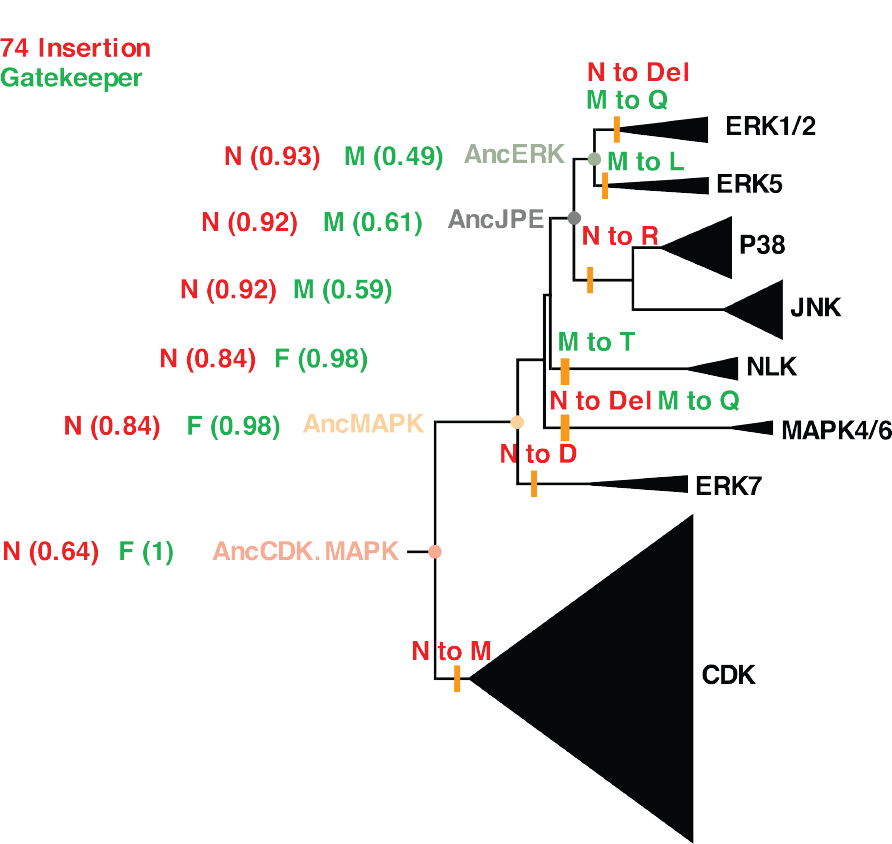
Phylogenetic tree indicating the mutations that occurred in the evolution from the common ancestor of CDK and MAPK to modern kinases. The status of the insertion (red) and identity of the gatekeeper residue (green) are indicated in the context of the CDK/MAPK group phylogenetic tree. The positions of ancestral nodes analyzed in this study are indicated by colored circles. Numbers in the brackets indicate the posterior probability for ancestral reconstructions. All posterior probabilities for the most recent evolutionary mutations are > 0.99.

**Figure 4-figure supplement 1:**
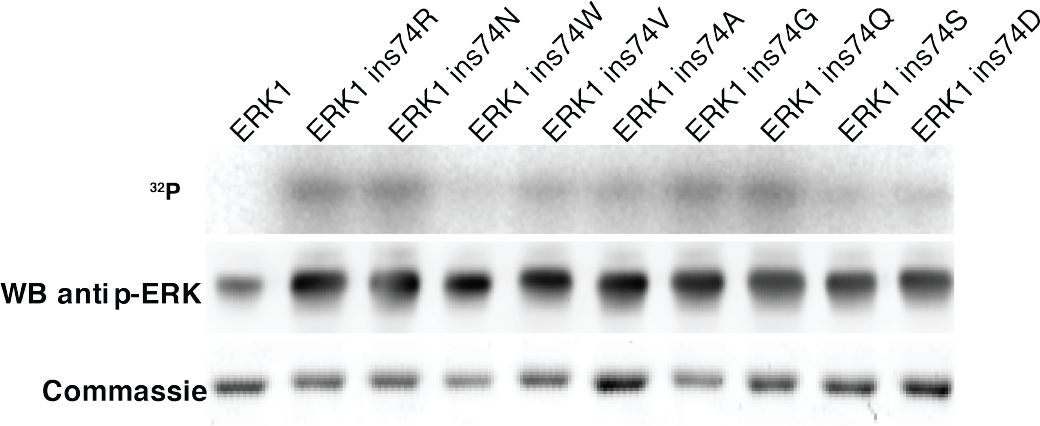
Other amino acid insertions after ERK1 residue 74 also increase activity. Kinase assays measuring the activity of different amino acids insertions after residue 74 in the β3-αC loop (top), Western blot analysis of ‘TEY’ phosphorylation (middle). and Coomassie brilliant blue staining loading control (bottom).

**Figure 4-figure supplement 2:**
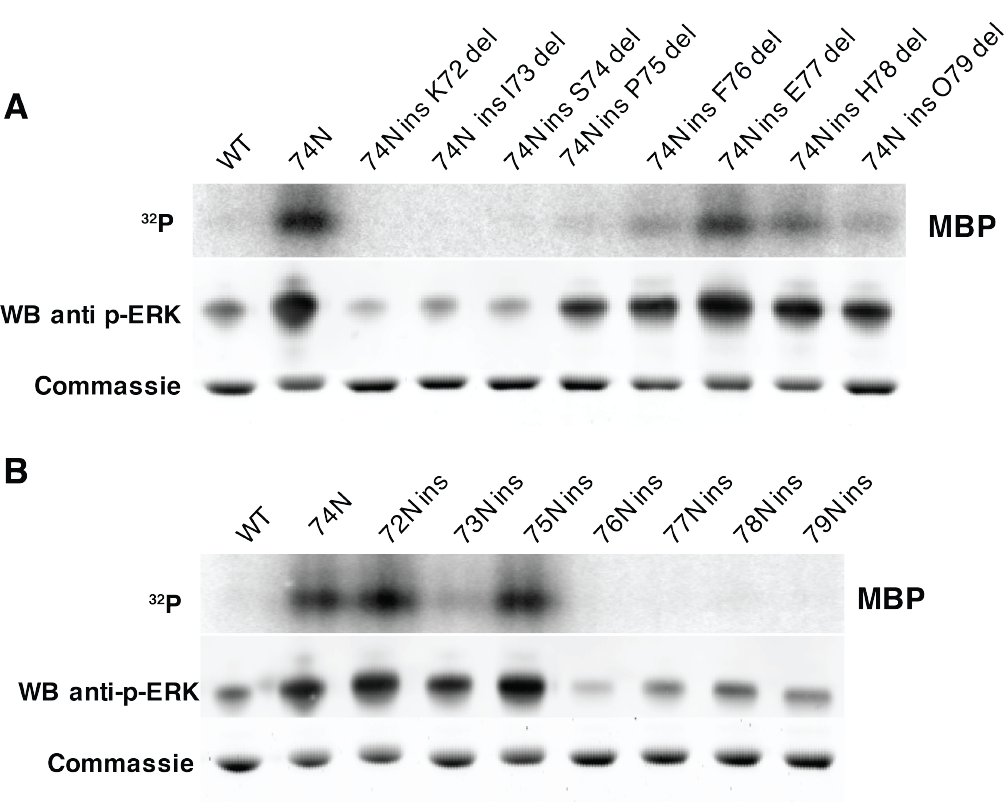
The length of the linker loop between K71 and P75 determines kinase activity. (A) Kinase assays measuring the activity of several mutants, in which one residue was deleted before or after Erk1 ins74N in β3-αC loop (top), Western blot analysis of ‘TEY’ motif phosphorylation (middle), and Coomassie brilliant blue loading control (bottom) (B) Kinase assays measuring the activity of mutants with asparagine insertion at multiple positions in the β3-αC loop (top), Western blot analysis of ‘TEY’ motif phosphorylation (middle), and Coomassie brilliant blue loading control (bottom).

**Figure 4-figure supplement 3:**
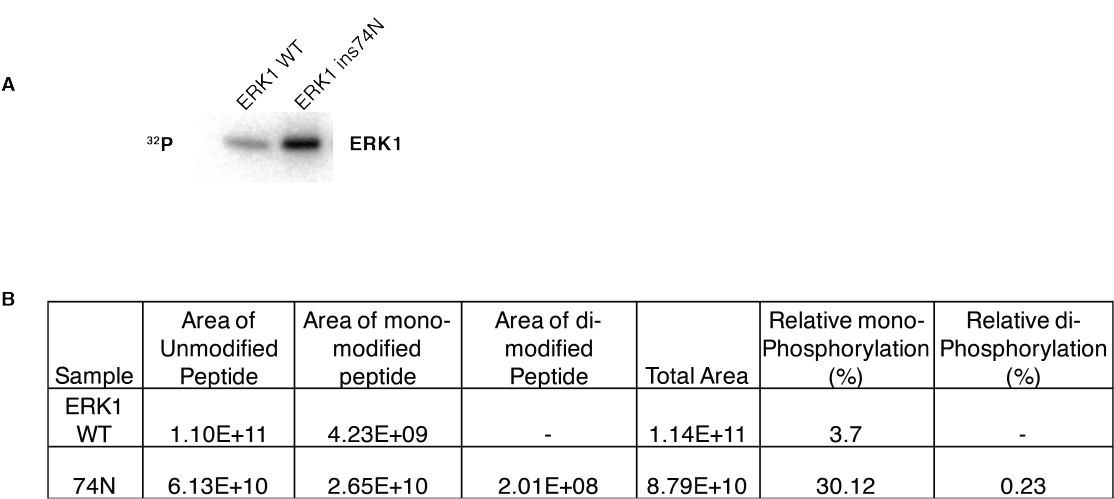
The ERK1 ins74N mutant has increased autophosphorylation on the TEY motif. (A) Autophosphorylation assay: purified ERK1 and ERK1 ins74N were incubated with kinase assay buffer for 30 minutes without substrate. (B) LC/MS analysis of the phosphorylation level of the ‘TEY’ motif. The area of the mass spectrometry peak corresponding to peptides containing ‘TEY’ were quantified. Mono-phosphorylated peptides include the peptides in which either threonine or tyrosine was phosphorylated. Di-phosphorylated peptides include only the peptides in which both threonine and tyrosine were phosphorylated. (-) indicates that no peptide was detected.

**Figure 5-figure supplement 1:**
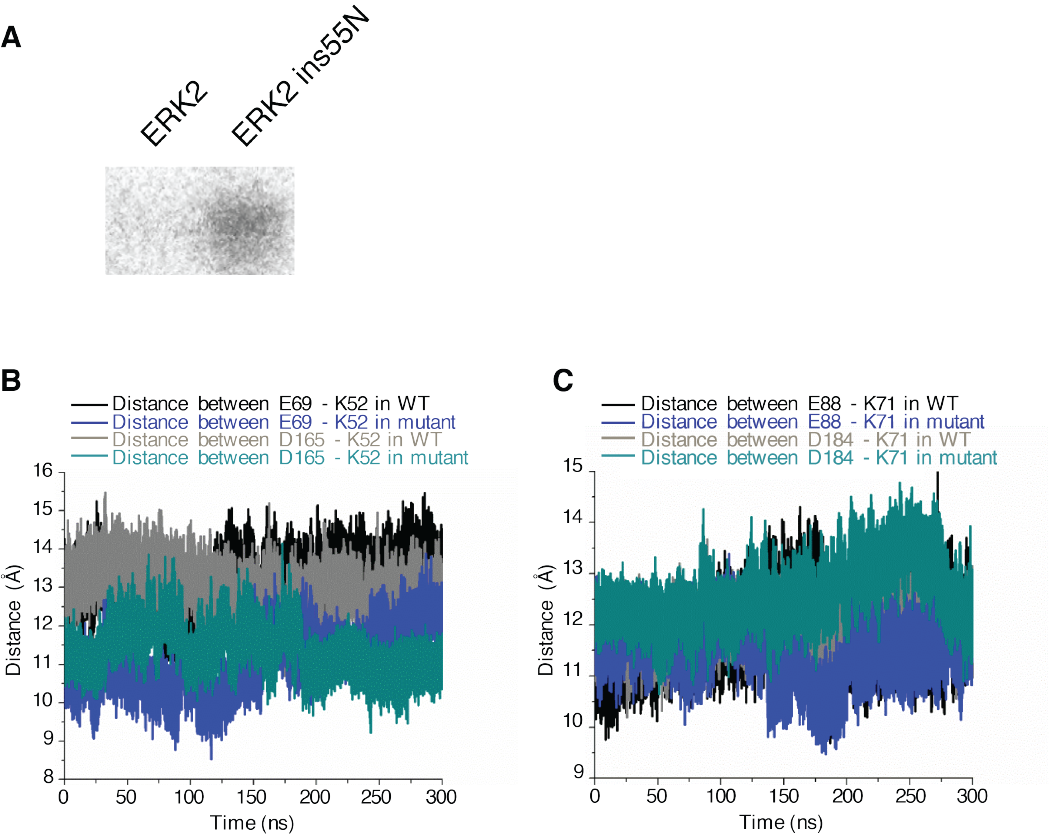
Molecular Dynamics simulation timecourse. (A) Kinase assays comparing the phosphorylation of MBP by ERK2 and ERK2 ins55N. (B − C) Distances from the C helix glutamate to the β3 strand lysine (ERK2 and ERK1, black; insertion mutants, blue); and from the β3 lysine to the DFG aspartate (ERK2 and ERK1, grey; insertion mutants, green) during 300 ns of simulation.

